# Electrostatic interactions at the five-fold axis alter heparin-binding phenotype and drive EV-A71 virulence in mice

**DOI:** 10.1101/648253

**Authors:** Han Kang Tee, Chee Wah Tan, Thinesshwary Yogarajah, Michelle Hui Pheng Lee, Hann Juang Chai, Nur Aziah Hanapi, Siti R. Yusof, Kien Chai Ong, Vannajan Sanghiran Lee, I-Ching Sam, Yoke Fun Chan

## Abstract

Enterovirus A71 (EV-A71) causes hand, foot and mouth disease epidemics with neurological complications and fatalities. However, the neuropathogenesis of EV-A71 remains poorly understood. In mice, adaptation and virulence determinants have been mapped to mutations at VP1-145, VP1-244 and VP2-149. We hypothesized that heparin-binding phenotype shapes EV-A71 virulence in mice. We constructed six viruses with varying residues at VP1-98, VP1-145 (which are both heparin-binding determinants) and VP2-149 (based on the wild type 98E/145Q/149K, termed EQK) to generate KQK, KEK, EEK, EEI and EQI variants. We demonstrated that the weak heparin-binder EEI was highly lethal in mice. The initially strong heparin-binding EQI variant acquired an additional mutation VP1-K244E, which confers weak heparin-binding phenotype resulting in elevated viremia and increased brain inflammation and virus antigens in mice, with subsequent high virulence. EEI and EQI-K244E variants inoculated into mice disseminated efficiently and displayed high viremia. Increasing polymerase fidelity and impairing recombination of EQI attenuated virulence, suggesting the importance of population diversity in EV-A71 pathogenesis *in vivo*. Combining *in silico* docking and deep sequencing approaches, we inferred that virus population diversity is shaped by electrostatic interactions at the five-fold axis of the virus surface. Electrostatic surface charges facilitate virus adaptation by generating poor heparin-binding variants for better *in vivo* dissemination in mice, likely due to reduced adsorption to heparin-rich peripheral tissues, which ultimately results in increased neurovirulence. The dynamic switching from heparin-binding to weak heparin-binding phenotype *in vivo* explained the neurovirulence of EV-A71.

**Author summary:** Enterovirus A71 (EV-A71) is the primary cause of hand, foot and mouth disease, and it can also infect the central nervous system and cause fatal outbreaks in young children. EV-A71 pathogenesis remains elusive. In this study, we demonstrated that EV-A71 variants with strong affinity to heparan sulfate (heparin) have a growth advantage in tissue culture, but are disadvantageous *in vivo*. When inoculated into mice, strong heparin-binding virus variants are more likely to be adsorbed to peripheral tissues, resulting in impaired ability to disseminate and being cleared from the bloodstream rapidly. The lower viremia level resulted in no neuroinvasion. In contrast, weak heparin-binding variants show greater levels of viremia, dissemination and subsequent neurovirulence in mice. We also provide evidence that the ability of EV-A71 to bind heparin is mediated by electrostatic surface charges due to amino acids on the virus capsid surface. In mice, EV-A71 undergoes adaptive mutation to acquire greater negative surface charges, thus generating new virulent variants with weak heparin-binding which allows greater viral spread. Our study underlines the importance of electrostatic surface charges in shaping EV-A71 virulence.

## Introduction

Enterovirus A71 (EV-A71) causes cyclical outbreaks of hand, foot and mouth disease (HFMD) in the Asia-Pacific region [1]. HFMD primarily affects children younger than 5 years old. The clinical manifestations are usually mild and characterized by fever, oral ulcers and skin rashes on hands and feet [2, 3]. In some cases, infection also results in severe neurological complications, including encephalitis, aseptic meningitis, acute flaccid paralysis and death [4]. There are no licensed antivirals, and licensed vaccines are only available in China. The EV-A71 genome encodes a polyprotein with a single open reading frame (ORF) flanked by 5’ and 3’ untranslated region [5]. The polyprotein is cleaved into four capsid proteins (VP1 to VP4) and seven nonstructural proteins (2A, 2B, 2C and 3A to 3D). The capsids form a protomer, and five protomers form a pentamer, and then twelve pentamers assembled around a genome forming a provirion. The five-fold axis symmetry is formed by VP1 and surrounded by a canyon. Virus-receptor binding at the canyon or other physical alterations such as heat, will displace the lipid pocket factor, and triggers viral uncoating [6].

Multiple receptors including human scavenger receptor class B2 (SCARB2) [7], P-selectin glycoprotein ligand-1 (PSGL-1) [8], heparan sulfate [9], sialylated glycan [10, 11], annexin II [12], vimentin [13] and nucleolin [14] have roles in EV-A71 attachment or entry. Following infection, EV-A71 can disseminate to different organs and invade the human central nervous system (CNS) [15, 16]. However, the neuropathogenesis of EV-A71 remains unclear and virulence determinants are not well elucidated. Recent reports have shown that EV-A71 strains with VP1-G/Q145 were more frequently isolated from severe HFMD with neurological complications [17–21]. However, the results contradicted *in vivo* studies which showed that VP1-E145 strains but not Q/G145 exhibit higher virulence and lethality in murine models [22–25]. These findings were also consistent with studies in cynomolgus monkeys, in which strong selection of VP1-E145 over VP1-G145 was observed [26]. VP1-Q/G145 residues but not VP1-E145 are responsible for binding to the receptor PSGL-1 [27]. Both VP1-E145 and VP2-I149 mutations conferred murine cell adaptation and increased mouse virulence [22, 24].

Neurotropism and neurovirulence has been previously related to the virus binding to heparan sulfate (heparin), a negatively charged glycosaminoglycan (GAG) found abundantly on most cell surfaces. Strong affinity for heparin was reported to cause attenuation in viruses such as Theiler’s murine encephalomyelitis virus [28], Japanese encephalitis virus [29], Murray Valley encephalitis virus [29], West Nile virus [30], yellow fever virus [31] and tick-borne encephalitis virus [32]. In contrast, virus variants with weak heparin-binding resulted in higher mortality in mice, as shown for Sindbis virus [33] and eastern equine encephalitis virus (EEEV) [34]. Interestingly, variants with strong heparin-binding can contribute to a higher neurovirulence if inoculated directly into the CNS, as demonstrated with EEEV [35].

EV-A71 also utilizes heparan sulfate (heparin) as an attachment receptor [9]. Amino acids clustered around the five-fold symmetry axis, specifically VP1-98, VP1-145, VP1-242 and VP1-244 modulate positive charges required for heparin-binding [36]. Positively-charged VP1-Q/G145 residues are associated with stronger heparin-binding, but not negatively-charged VP1-E145. The propensity of EV-A71 to acquire positively-charged residues at the five-fold axis is associated with increased heparin-binding upon *in vitro* culture adaptation [36]. In contrast, experimental studies of VP1-E145 in mice and monkey studies showed higher virulence and lethality, and the role of heparin-binding *in vivo* has not been fully characterized.

In the current study, we sought to delineate the discrepancy exhibited between *in vitro* cytopathogenicity due to heparin-binding and association of weak heparin-binding with *in vivo* virulence. We demonstrated that acquisition by EV-A71 of negative charges at the five-fold axis reduced heparin-binding capacity. This resulted in high viremia and enhanced lethality in mice.

## Results

### Construction and rescue of clone-derived virus variants

We have previously described the role of VP1-98 and VP1-145 as modulators of heparin-binding in cell culture [36]. As the role of heparin remains unknown *in vivo*, in the present study, we elucidated the role of heparin-binding in shaping EV-A71 virulence in mice. Five EV-A71 variants were engineered using a wild type EV-A71 strain (5865/SIN/00009, subgenogroup B4), which possess VP1-E98, VP1-Q145 and VP2-K149 residues (designated as EQK following amino acid residue order). Site-directed mutagenesis was performed to generate different combinations of variants, denoted as KQK, KEK, EEK, EEI and EQI (Fig 1A). Some variants had the mouse-adaptive determinant, K149I [22, 24, 37] along with substitutions in the heparin-binding determinants, VP1-98 and VP1-145. All EV-A71 variants were viable with comparable plaque morphology (Fig 1B) and were confirmed as genetically stable by sequencing (data not shown).

**Fig 1.**
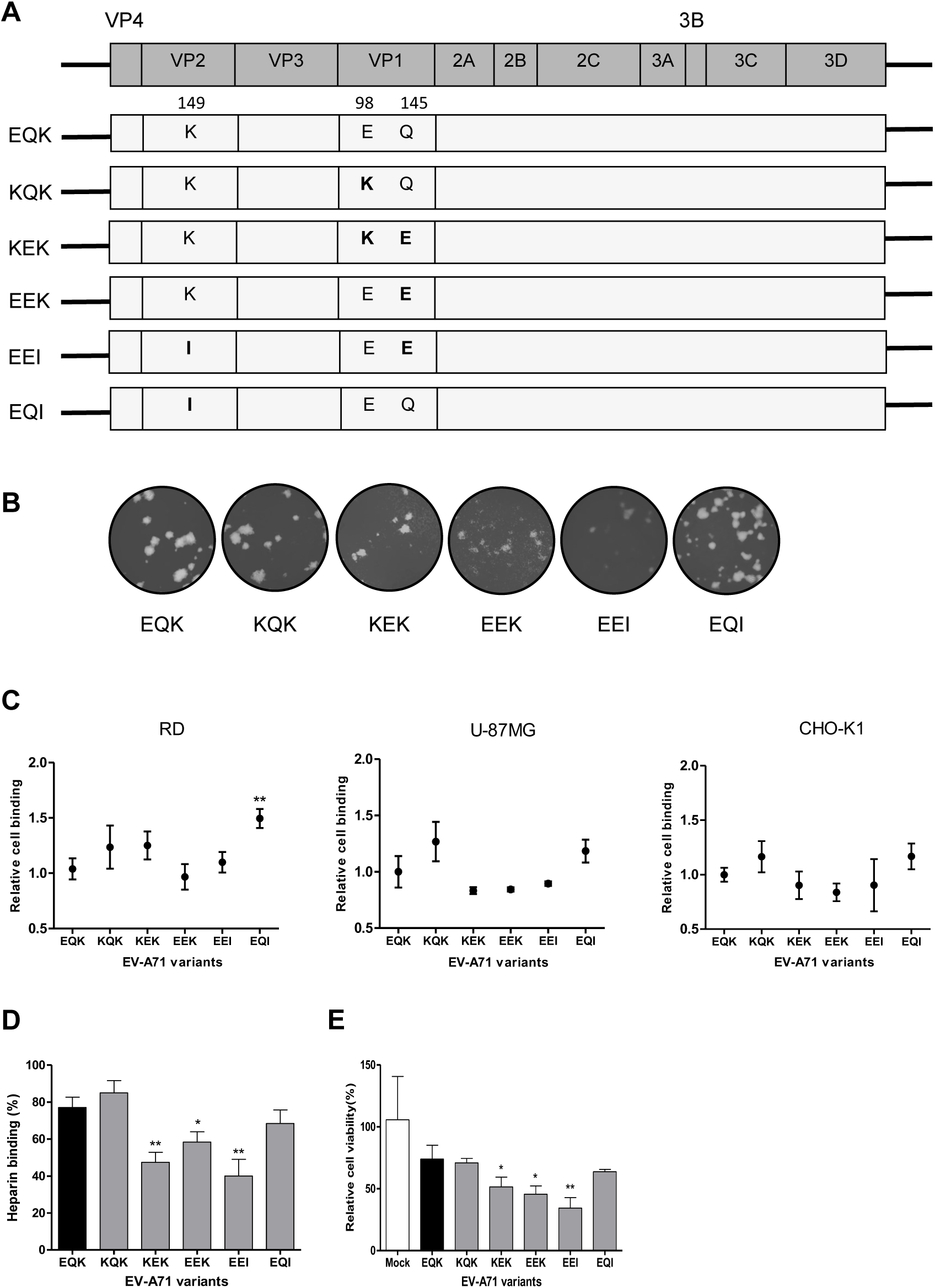
*In vitro* characterization of constructed EV-A71 variants. (A) Schematic illustration of the EV-A71 genome and the infectious clone constructs. Different amino acids were substituted at VP1-98, VP1-145 and VP2-149 sites (labeled in bold), with reference to the wild type strain EQK. (B) The clone-derived EV-A71 variants were propagated in RD cells and showed comparable plaque morphologies. (C) Relative cell binding of EV-A71 variants were measured using a virus attachment assay in RD, U-87MG and CHO-K1 cells. The relative cell binding value was determined by absorbance ratio of each variant over WT virus. (D) The binding affinity of EV-A71 variants to heparin sepharose beads was analyzed. (E) Inhibitory effect of heparin on EV-A71 variants was evaluated by pretreating the viruses with soluble heparin before infection of RD cells. Results are presented as mean ± SD (n=3). Error bars indicate standard deviations from triplicates. Statistical significances are denoted with \**P* < 0.05, \*\**P* < 0.01 as compared to the WT.

### VP1-145 modulates heparin-binding phenotypes of EV-A71 in *in vitro* cell culture

Next, we analyzed the cell-binding ability of EV-A71 variants in RD (human muscle cells), U-87MG (human neuronal cells) and CHO-K1 (murine cells) cells. In general, all six variants showed consistent binding phenotypes across human (RD) and murine (CHO-K1) cells (Fig 1C). There was also no significant difference in receptor binding between RD and U-87MG cells, suggesting no preferential virus tropism between human muscle and neuronal cell lines. Consistent with our previous findings [36], EV-A71 variants with VP1-E145 (K<u>E</u>K, E<u>E</u>K and E<u>E</u>I) exhibited lower binding capacity compared to those with VP1-Q145 (E<u>Q</u>K, K<u>Q</u>K, E<u>Q</u>I). With the additive effect of positively-charged VP1-K98, KQK showed a higher binding phenotype than wild type EQK across all the cell lines. However, the higher binding phenotype exhibited by KEK in RD cells compared to wild type could be linked to the compensatory mutation we reported previously in which VP1-K98 restored virus binding ability in the presence of VP1-E145 [36].

We further investigated the heparin-binding ability of the EV-A71 variants. EV-A71 variants with VP1-E145 (KEK, EEK and EEI) displayed significant reduction of 29.7%, 18.8% and 37.1% in heparin-binding, respectively (Fig 1D). Similar findings were also observed in a heparin inhibition assay. Pre-treated heparin inhibition of these three VP1-E145 variants significantly reduced RD cell viability to 51.5%, 45.5% and 34.3%, respectively (Fig 1E). Based on these heparin-binding results, the EV-A71 variants were categorized into two groups: strong heparin binder (EQK, KQK and EQI) and weak heparin binder (KEK, EEK and EEI).

### Potential association of weak heparin-binding variants with *in vivo* virulence

To determine the role of heparin-binding in EV-A71 *in vivo* virulence, we performed intraperitoneal (i.p.) infection of one-day old suckling mice with 1 × 10^5^ PFU of each EV-A71 variant. The clinical score and survival analysis of infected mice are shown in Fig 2A. EEI-infected mice exhibited the highest virulence *in vivo*, with all the infected mice dying by day 4 post-infection. Notably, 66.7% of the EQI-infected mice died by day 12 post-infection, followed by 20% mortality in EEK-infected mice. Consistent with previous findings [22–24, 38], EV-A71 variants with VP1-E145 were associated with an increased virulence phenotype in animal models, except for KEK. Mice infected with other EV-A71 variants showed no apparent clinical signs and survived beyond 13 days post-infection. These data suggest that *in vivo* pathogenicity in mice was increased by an additional VP2-K149I mutation and further enhanced by weaker heparin-binding conferred by VP1-E145, as demonstrated in the EEI-infected mice.

**Fig 2.**
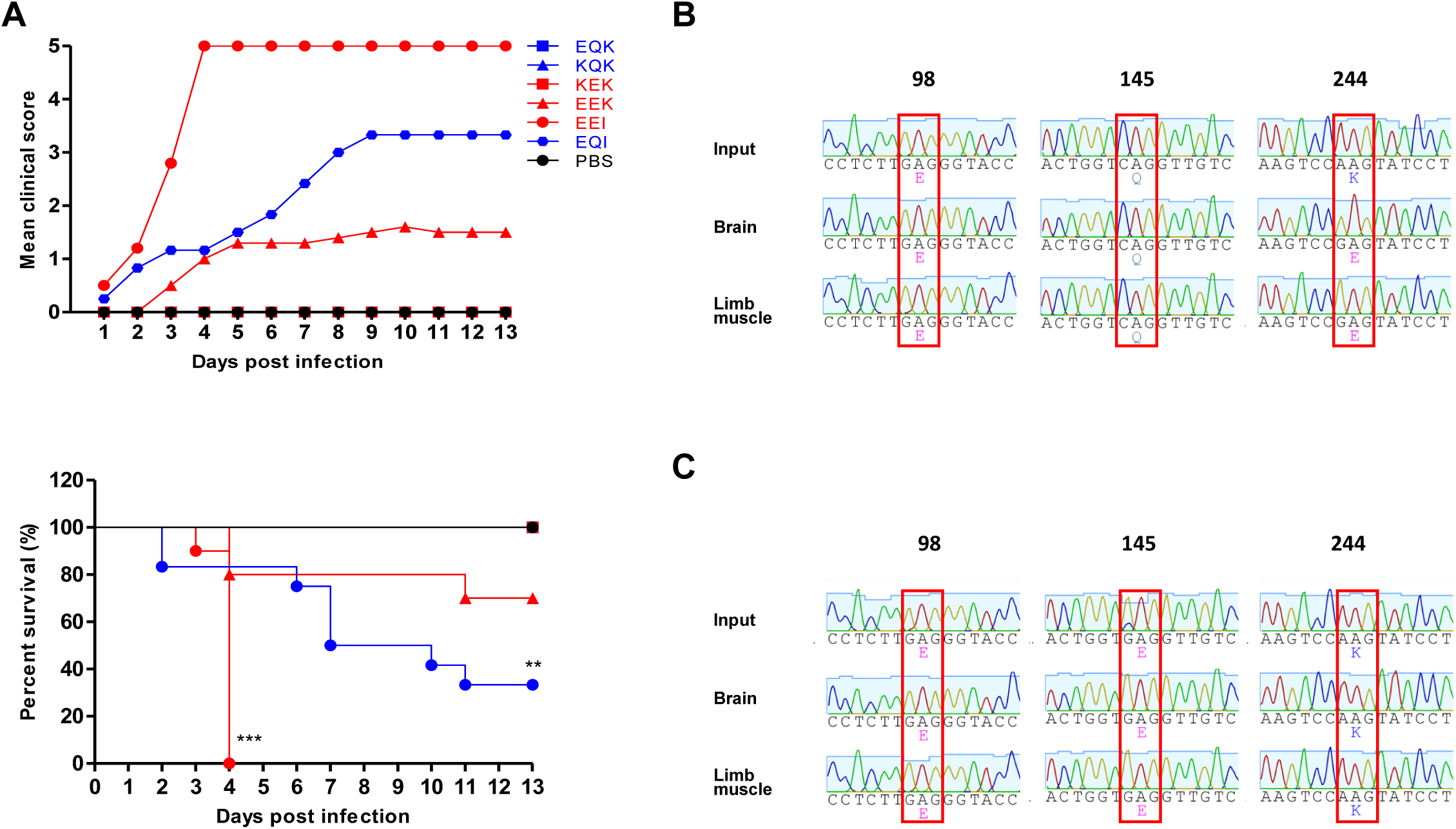
Clinical scores and survival rates of suckling mice infected with EV-A71 variants. One-day old suckling mice (n=9-12) were inoculated with 1 × 10^5^ PFU of different EV-A71 variants by i.p. injection. (A) A litter of mock-infected mice was used as a control group receiving PBS injection. The clinical scores and percentage of survival of the infected mice groups were monitored daily for 13 days. The severity of clinical symptoms was scored as follows: 0, healthy; 1, weak or less active; 2 hunched posture and lethargy; 3, one-limb paralysis; 4, two-limb paralysis; 5, moribund or dead. Significant differences compared to EQK are labelled as * (*P* < 0.05) and \*\*\**P* (< 0.001). EQK, KQK and KEK curves are identical to that of the mock-infected group. The VP1 sequence chromatograms of EQI (B) and EEI (C) populations isolated from infected animal organs are shown, highlighting VP1-98, VP1-145 and VP1-244 (note that VP2-149 is not shown). The emergence of E244 virus isolated from brains and limbs of EQI-infected moribund mice are shown.

We further asked why the EQI variant, a strong heparin binder, exhibited a relatively high virulence in mice. Viral genomic RNA from the brains and hind limbs of dead EQI-infected mice were harvested for genome sequencing, revealing that the EQI variant had acquired a VP1-K244E mutation (Fig 2B). This mutation, however, was not present in the EEI-infected mice (Fig 2C).

### Emergence of EQI-K244E variant resulted in abolished heparin-binding ability and regaining of virulence

Since VP1-K244 is a key determinant of heparin-binding [36], we speculated that the emergence of VP1-K244E had abolished the heparin-binding ability of the EQI variant. To investigate the role of VP1-K244E mutation in heparin-binding and virulence in mice, we introduced this mutation into the EQI variant through site-directed mutagenesis. The EQI variant with K244E mutation (termed EQI-K244E) however failed to achieve high virus yield in tissue culture for subsequent *in vivo* experiments (data not shown). We thus collected EQI-K244E and EEI from the brain homogenates (indicated with ^+^) for subsequent experiments, after confirming the sequences using Sanger sequencing. EQI-K244E^+^ displayed significant reduction of heparin-binding compared to clone-derived EQI and EEI, and EEI^+^ (Fig 3A). To determine the association of heparin-binding and *in vivo* virulence, we then infected one-day old suckling mice with clone-derived EEI and EQI-K244E^+^ by i.p. administration. Brain homogenate from EQI-infected surviving mice was also harvested and used as a negative control (EQI^+^; viral RNA not detected in RT-PCR). At day 4 post-infection, 100% mortality was observed in EEI and EQI-K244E^+^-infected mice but none of the mice succumbed to EQI^+^ infection (Fig 3B).

**Fig 3.**
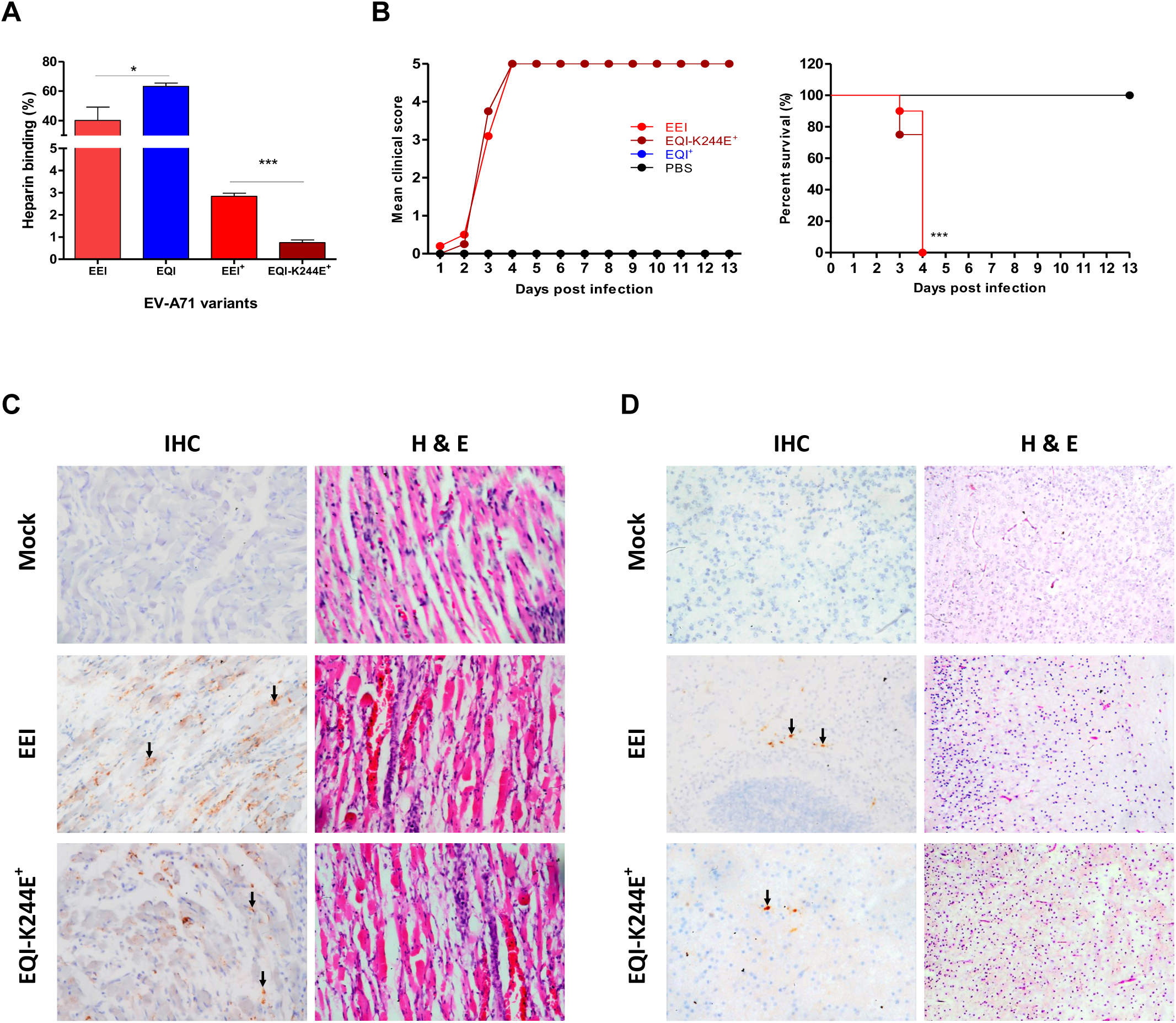
Characterization of virulent phenotype of EEI and EQI-K244E^+^ variants. (A) EV-A71 variants collected from the brain homogenates of dead infected mice were compared to clone-derived variants for heparin-binding properties. (B) One-day old suckling mice (n=8-12) were infected with EEI, EQI-K244E^+^ and EQI^+^ through i.p. injection. A control group receiving PBS injection was also included. The clinical scores and percentage of survival are shown over 13 days post-infection. Significant differences compared to WT are labelled as ** (*P* < 0.01) and \*\*\**P* (< 0.001). Tissue samples of mice which succumbed to EEI and EQI-K244E^+^ infection were subjected to IHC and H&E staining. Virus antigen was seen in muscle cells (C), along with increased inflammatory infiltrates. In brains (D), antigen-positive neurons were seen in the midbrain (representative image from an EQI-K244E^+^-infected brain) and pons (representative image from an EEI-infected brain), with severe inflammation in the cortex shown by H&E staining. Magnification for IHC staining: X40; H&E staining: X20.

The hind limb and brain samples from EEI and EQI-K244E^+^-infected mice were then processed for histopathological analysis, and results supported earlier findings. Immunohistochemical (IHC) examination revealed massive localization of viral antigens in both EEI and EQI-K244E^+^-infected muscles, indicating that skeletal muscle is an important replication site (Fig 3C). Inflammation and extensive muscle damage were also observed in the haematoxylin and eosin (H&E)-stained sections of muscle. In addition, viral antigens were detected in neurons mainly distributed in the pons and midbrains (Fig 3D). Mononuclear cells infiltrations were also evident in the cortices. In contrast, no distinctive histopathological change was observed in the mock-infected organ samples.

### High lethality of weak heparin-binding EV-A71 variants correlates with high viremia

Strong heparin-binding ability confers the advantage of promoting virus attachment on the cell surface, thus increasing the probability of virus-functional receptor interaction *in vitro* [9]. However, we have demonstrated that a strong heparin-binding phenotype is deleterious to virus pathogenesis *in vivo*. To unravel the discrepancy of cytopathogenicity *in vitro* and *in vivo* virulence, we investigated virus dissemination in mice. Following i.p. infection, five mice were sacrificed for viral load quantitation in brain and hind limbs. EEI and EQI-K244E^+^ variants replicated to higher titers than EQI in both hind limbs and brain, at 2 and 4 days post-infection (Fig 4A).

**Fig 4.**
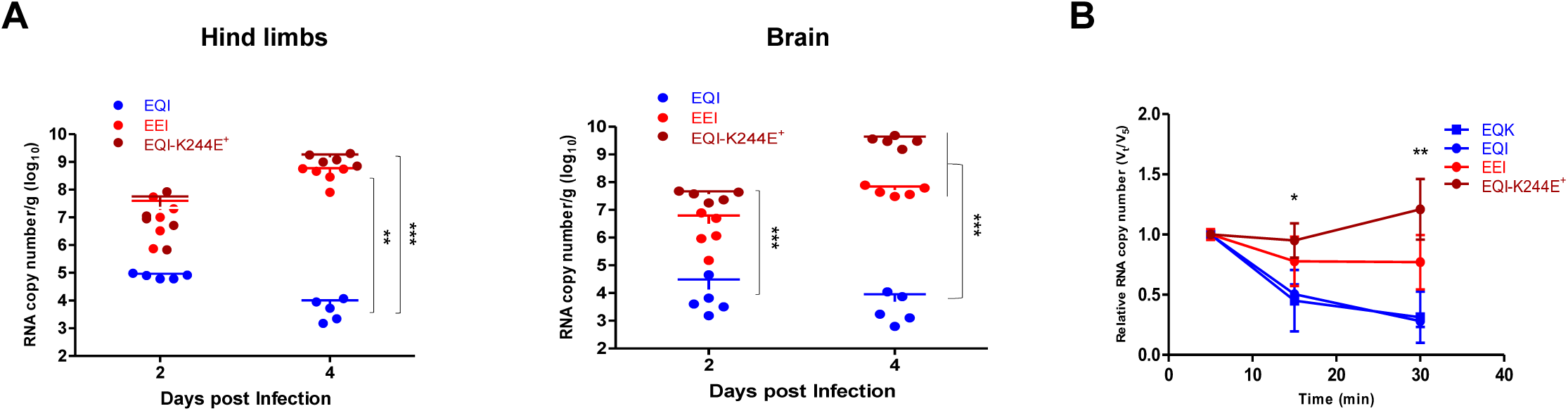
Viral load quantitation from harvested organs and viremia level induced by EV-A71 variants. (A)At selected time points, EQI, EEI and EQI-K244E^+^-infected mice (n=5) were sacrificed and viral loads were determined from harvested hind limbs and brains using qRT-PCR. Significant differences between viral variants are labelled as ** (*P* < 0.01) and *** (*P* < 0.001). (B) Virus clearance from blood was quantitated using qRT-PCR following intravenous inoculation of EQK (WT), EEI, EQI or EQI-K244E^+^ into 3-4 week-old mice. The virus titers were quantitated at selected timepoints up to 30 minutes. Significant differences between EQK and EQI-K244E^+^ are labelled as * (*P* < 0.05) and ** (*P* < 0.01).

To investigate the attribution of viremia to *in vivo* pathogenesis, three- to four-week-old mice were infected intravenously with EQK, EEI, EQI and EQI-K244E^+^ variants. Blood samples were then collected at 5, 15 and 30 min post-inoculation for viral load quantitation. Both strong heparin-binding variants, EQK and EQI showed rapid viral clearance, with approximately 70% cleared from the bloodstream at 30 minutes (Fig 4B). Only about 23% of EEI had been cleared by 30 minutes post-infection. A sustained viremia level with minimal clearance was observed in EQI-K244E^+^-infected mice. Our data indicated stronger heparin-binding resulted in low viremia levels in the host, which may be due to adsorption and sequestration of viruses in surrounding tissues. The low viremia level may render the virus less efficient in disseminating to other organs, as seen in EQK and EQI variants.

### High fidelity and impairing recombination attenuate *in vivo* virulence of EQI

The emergence of a weak heparin-binding variant with VP1-K244E mutation is the key determinant of *in vivo* adaptation and pathogenesis of EQI. We hypothesized that EQI is avirulent without the acquisition of the VP1-K244E mutation *in vivo*. To reduce mutation rates and restrict generation of viral quasispecies, we engineered the viral RNA-dependent RNA polymerase (RdRp) of EQI to harbor previously identified high-fidelity mutations G64R and L123F (abbreviated as HF in Fig 5A) [39–41] and recombination deficient mutation Y276H (labelled as Rec^-^) [42]. We employed a luciferase-based replicon system to assess the impact of these mutations on genome replication. As demonstrated in Fig 5B, no significant differences in luciferase activities were observed between wild type EV-A71 Nluc Rep, EV-A71 Nluc Rep-HF and EV-A71 Nluc Rep-Rec^-^, suggesting that these mutated RdRp variants replicate as efficiently as the wild type. Next, we generated and rescued the EQI-HF and EQI-Rec^-^ virus variants. These EQI-HF and EQI-Rec^-^ variants were genetically stable with no reversion of mutations and emergence of VP1-244E observed after a few passages, in addition to indistinguishable plaque morphology to EQI (data not shown).

**Fig 5.**
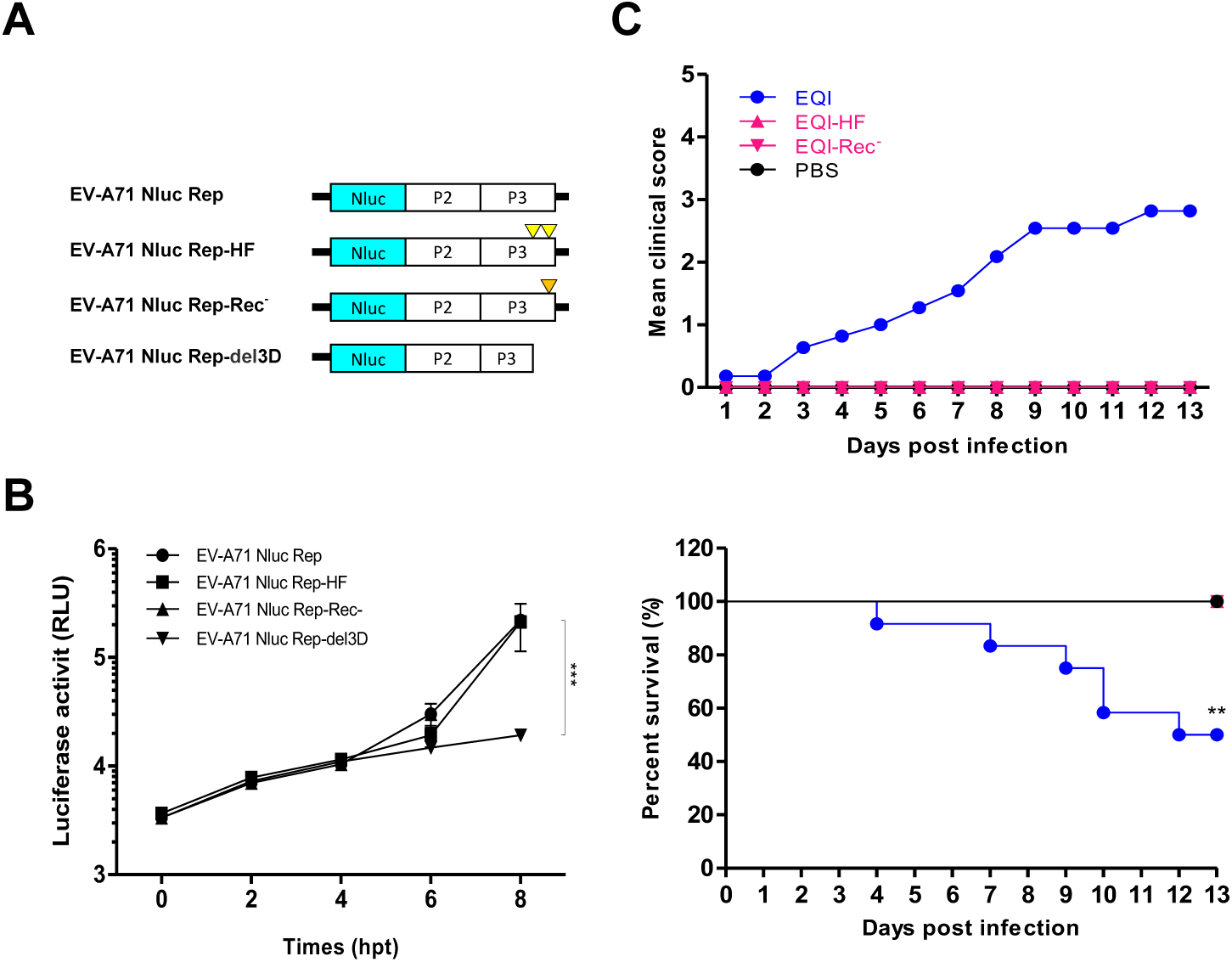
High fidelity and recombination deficiency restrict emergence of the VP1-K244E mutation, resulting in attenuation of EQI. The RdRp of EQI was modified to harbor the high fidelity G64R and L123F mutations (HF) or the recombination-deficient Y276H mutation (Rec^-^) and virulence was analyzed in mice. (A) EV-A71 subgenomic replicon (EV-A71 Nluc Rep), EV-A71 Nluc Rep-HF (HF mutations are indicated by yellow triangles), EV-A71 Nluc-Rep-Rec^-^ (Rec^-^ mutation is denoted by orange triangle) and truncated replicon (EV-A71 Nluc Rep-del3D) were transfected separately into RD cells. (B) Luciferase activities were determined up to 8 hours post-transfection. Significant differences between replicon variants and WT are labelled as *** (*P* < 0. 001). (C) Virulence of EQI-HF, EQI-Rec^-^ and EQI following i.p. infection of mice was measured by clinical scores and percentage of survival. EQI-HF and EQI-Rec^-^ curves are identical to that of the mock-infected PBS group. Significant difference between viral variants and EQI is labelled as ** (*P* < 0.01).

To characterize the impact of increased fidelity and recombination deficiency on *in vivo* virulence, one-day old suckling mice were infected with EQI, EQI-HF and EQI-Rec^-^. Half of the EQI-infected mice died by day 12 post-infection (Fig 5C), while none of the mice died following EQI-HF and EQI-Rec^-^ infection. A reduced ability of the EQI strain to undergo mutations was associated with loss of *in vivo* virulence, although we were unable to recover EQI-HF and EQI-Rec^-^ viruses from surviving mice to definitively show that this was due to lack of the VP1-K244E mutation.

### Emergence of VP1-K244E is important for neuroinvasion

To determine if emergence of the VP1-K244E mutation is critical for systemic dissemination, we next examined neurovirulence (the ability to directly infect the CNS) of all the EV-A71 variants following direct intracerebral inoculation. A dose of 1 × 10^5^ PFU of each of the EV-A71 variants was intracerebrally injected into separate litters of one-day old mice (Fig 6). Similar mortality rates (100% mortality at day 4 post-infection) were shown by the EEI and EQI-K244E^+^ variants, the former having been earlier shown to be highly lethal following i.p. infection (Fig 2A). Notably, EQI infection, which caused 66.7% mortality when inoculated intraperitoneally, now showed a remarkable drop of virulence to 8.3% mortality following i.c. infection. Detection of the VP1-K244E mutation by sequencing in organ samples from dead EQI-infected mice further confirmed its importance as a neuroinvasion determinant (data not shown). The remaining virus variants did not cause any disease symptoms in mice, and no virus was detected. Taken together, our data indicates that the critical mutation conferring neuroinvasive phenotype to the EQI variant is VP1-K244E, which mainly arises during systemic dissemination when EQI is inoculated intraperitoneally and not directly into the brain.

**Fig 6.**
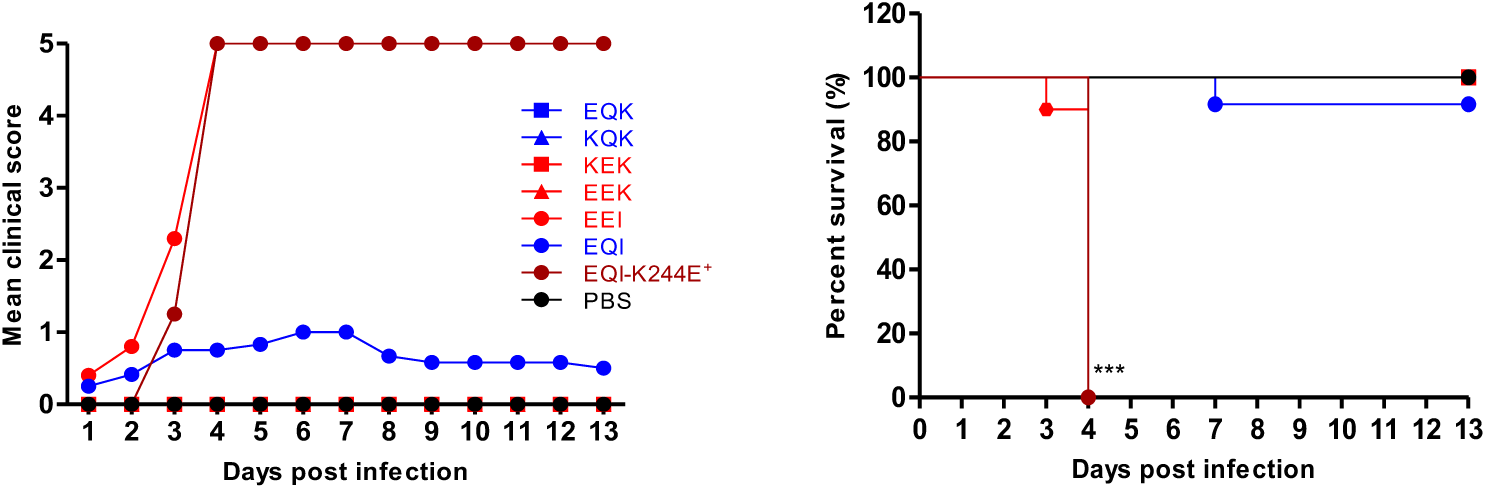
EQI virulence was attenuated when inoculated intracerebrally. A dose of 1 × 10^5^ PFU of each EV-A71 variant was administered intracerebrally into one-day old suckling mice. Clinical scores and percentage of survival over 13 days are shown. Significant differences between viral variants and WT are labelled as *** (*P* < 0.001).

### Uncharged VP1-244 intermediate variant emerged during transition to EQI-K244E

To investigate how the VP1-K244E mutation emerges during *in vivo* infection, a litter of 14 suckling mice was intraperitoneally infected with EQI. The mice were sacrificed at day 3, 7 and 11 post-infection or when moribund to harvest hind limbs and brains for next-generation sequencing of the virus population diversity. At day 9 post-infection, two moribund mice were collected. We first screened all the harvested samples using RT-PCR. At day 3 post-infection, none of the collected organs were positive for EV-A71 (Fig 7A). All five muscle samples collected were positive for EV-A71 at day 7 post-infection, suggested that viruses were replicating in skeletal muscles, but brain samples were negative suggesting that limited virus had disseminated to the brain at this time point. As expected, both muscle and brain samples collected from the moribund mice at day 9 post-infection were positive for EV-A71. None of the remaining mice collected at day 11 post-infection were positive for EV-A71 RNA in both hind limbs and brain.

**Fig 7.**
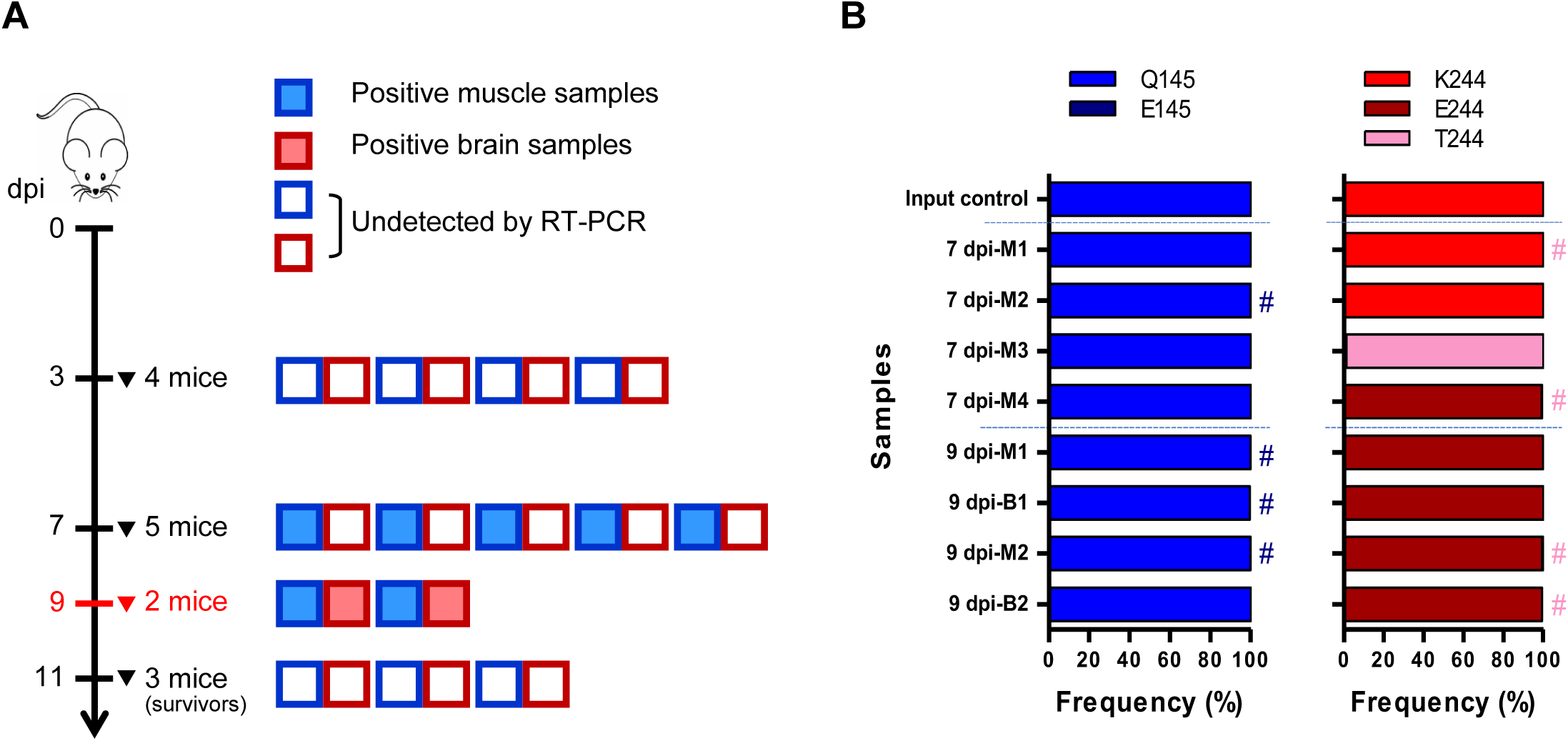
Sequential emergence of K244E mutation involves intermediate transition variants. One-day old suckling mice were intraperitoneally injected with 1 × 10^5^ PFU of EQI. (A) At days 3 and 7 post-infection, mice were euthanized to harvest their hind limb muscles and brains. At day 9 post-infection, two moribund mice were sacrificed for processing (highlighted in red). At the end of the experiment (day 11 post-infection), the remaining three healthy mice were sacrificed and categorized as ‘survivors’. The boxes indicate RT-PCR results for EV-A71 for each muscle and brain sample. A total of eight samples were positive, comprising four muscle samples from day 7 post-infection, and two muscle and brain samples from day 9 post-infection. (B) NGS results showing the frequency of different variants at the VP1-145 and VP1-244 sites. Variant frequencies lower than 0.1% are indicated with #. Noted that 7 dpi-M5 sample was excluded from analysis due to poor sequencing coverage detected.

Deep sequencing of the EV-A71-positive organ samples revealed that the VP1-Q145 residue was highly stable with > 99% variant frequency (Fig 7B). The frequency of VP1-Q145E mutation in these samples was lower than 0.1% (labelled with # in Fig 7B). To our surprise, 7 dpi-M5 sample displayed a poor sequencing coverage and therefore was eliminated from the analysis. We observed a sequential transition of K244 to E244 from day 7 to day 9 post-infection. At day 7 post-infection, VP1-K244 was predominant in two out of four hind limb samples (M1 and M2), while only a single sample (M4) showed VP1-E244 as the dominant viral population. Notably, we identified a novel substitution, VP1-K244T from the hind limb of sample M3 with high frequency of 90%. As infection progressed, VP1-E244 was solely detected in the organs harvested at day 9 post-infection, further affirming the contribution of VP1-E244 to *in vivo* virulence. Apart from VP1-244 and 145, we also detected other non-synonymous mutations previously linked to heparin-binding from the harvested samples, including VP1-L97R, N104S and E167G (Table 1). The emergence of uncharged T244 intermediate during the transition from positively-charged K244 to negatively-charged E244 showed the high plasticity of EV-A71 in generating compensatory mutations *in vivo*.

**Table 1.** Non-synonymous mutations related to HS binding detected from different organ samples of EQI-infected mice

| Samples | L97 |  | E98 |  | N104 |  | Q145 |  | E167 |  | K244 |  |
| --- | --- | --- | --- | --- | --- | --- | --- | --- | --- | --- | --- | --- |
|  | Mutation | Frequency (%) | Mutation | Frequency (%) | Mutation | Frequency (%) | Mutation | Frequency (%) | Mutation | Frequency (%) | Mutation | Frequency (%) |
| **Input control (EQI)** |  |  | E98Q | 6.06 |  |  |  |  | E167G | 19.11 |  |  |
|  |  |  | E98G | 4.69 |  |  |  |  | E167Q | 3.84 |  |  |
|  |  |  | E98V | 2.32 |  |  |  |  |  |  |  |  |
| **7 dpi M1** |  |  |  |  |  |  |  |  |  |  |  |  |
| **7 dpi M2** |  |  |  |  |  |  |  |  | E167G | 99.77 |  |  |
| **7 dpi M3** |  |  |  |  | N104S | 99.34 |  |  |  |  | K244T | 98.71 |
| **7 dpi M4** |  |  |  |  | N104S | 98.25 |  |  |  |  | K244E | 99.16 |
| **9 dpi B1** |  |  |  |  | N104S | 99.88 |  |  |  |  | K244E | 99.83 |
| **9 dpi M1** |  |  |  |  | N104S | 99.45 |  |  |  |  | K244E | 99.86 |
| **9 dpi B2** |  |  |  |  |  |  |  |  |  |  | K244E | 99.12 |
| **9 dpi M2** | L97R | 99.49 |  |  |  |  |  |  |  |  | K244E | 99.45 |

\*Non-synonymous mutations are categorized into three different groups based on their frequency of detection in NGS. Variant frequencies of 1-90% and > 90% are highlighted in blue and green, respectively.

### Weak heparin-binding is due to loss of electrostatic interactions at the five-fold axis

EEI was experimentally proven to be highly lethal in mice. However, we observed that EQI selectively acquires VP1-K244E over the VP1-Q145E mutation to gain neuroinvasive and neurovirulent properties (Figs 3B & 6). We reasoned that VP1-E244 exhibits weaker heparin-binding ability compared to VP1-E145, and therefore, could be favorably selected *in vivo*. We employed *in silico* analysis to characterize the heparin-binding affinity of VP1-E145 and VP1-E244. VP1-98, 145 and 244 are located around the five-fold axis of the EV-A71 pentamer (Fig 8A). Based on the electrostatic maps, the five-fold axis of the EQI variant is highly positive-charged (Fig 8B), implying strong affinity to heparin. With the VP1-Q145E substitution, EEI has lower positive charges at its five-fold axis. Since the VP1-K244 residue is protruding from the surface of the five-fold axis, substitution of a positively-charged lysine residue to a negatively-charged glutamic acid at this position greatly reduces the electrostatic potential on the five-fold axis and changes the capsid conformation.

**Fig 8.**
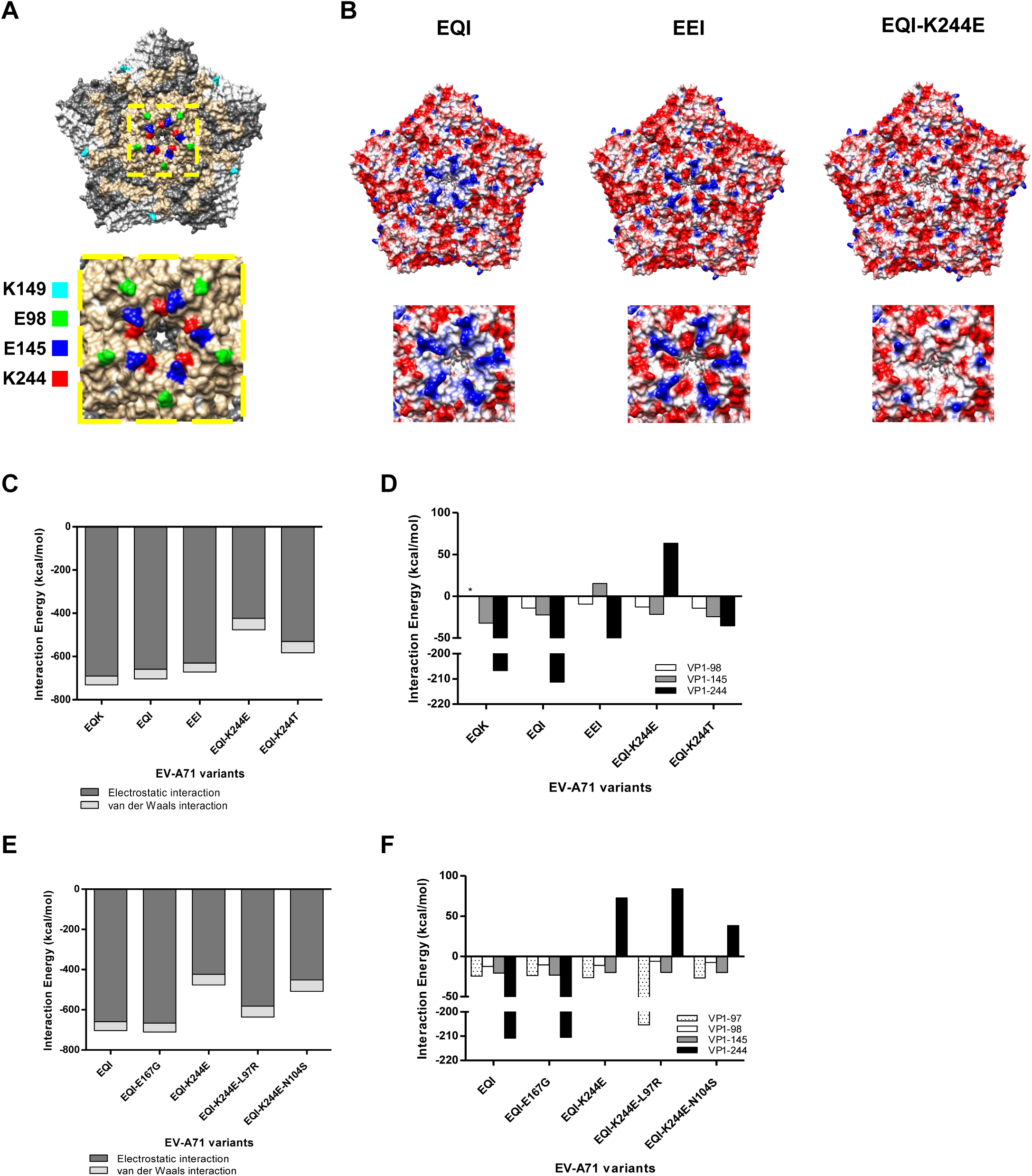
Comparison of electrostatic surface properties of the EV-A71 structure. The analyzed amino acid residues are labelled in different colors in the EV-A71 capsid pentamer structure (PDB ID: 4AED). (A) The front views of the EV-A71 pentamer are displayed along with a magnified view of the five-fold axis using Chimera 1.10.1. (B) Electrostatic surface properties of EQI, EEI and EQI-K244E variants were examined. Electrostatic charges are contoured from red (-5 kcal/mol·e, negatively-charged) to blue (+5 kcal/mol·e, positively-charged). Magnified views of the five-fold axis are shown for each variant. Total interaction energy (C) and individual interaction energy of residues (D) within a 4Å radius of EV-A71 variants and VP1-244 variants docked to heparin were evaluated. Total interaction energy (E) and individual interaction energy of residues (F) within a 4Å radius of other mutations detected from NGS docked to heparin were examined.

Simulated docking of 12-mer heparin to EV-A71 residues within a 4Å radius revealed no notable change in interaction energies between EQK (-730.76 kcal/mol) and EQI (-703.71 kcal/mol) (Fig 8C). Substitution of the VP1-Q145E mutation in EEI resulted in a drop of 32.34 kcal/mol interaction energy when compared to EQK. The EQI-K244E variant showed the weakest heparin-binding ability as the interaction energy drastically dropped to -476.32 kcal/mol. Compared to the EQI-K244E variant, uncharged EQI-K244T variants exhibited higher interaction energies of -552.21 kcal/mol and -582.89 kcal/mol, respectively. The energy change was mainly contributed by the VP1-244 residue, with a descending energy order of K244, T244 and E244, which correlates with the scale of strong to weak heparin-binding (Fig 8D).

When examining the heparin docking of non-synonymous variants detected by NGS, the VP1-E167G mutation had no effect on interaction energy compared to EQI (Fig 8E). Interestingly, variants with VP1-L97R and VP1-N104S mutations showed slight increase in the interaction energy of EQI-K244E variants, suggesting an enhanced effect of heparin-binding. Although EQI-K244E-L97R showed very weak binding strength at the VP1-K244E site similar to EQI-K244E, a compensatory effect was seen in L97R site which singly contributes to a more negative interaction energy i.e. strong heparin-binding (Fig 8F). Meanwhile, VP1-N104S mutation has also increased the heparin-binding of EQI-K244E variant. Taken together, different compensatory mutations emerged to overcome the capsid instability and alter the virus fitness.

## Discussion

As heparin-binding phenotype has been implicated in virulence of some neurotropic viruses, we studied the relationship between heparin-binding phenotype and mouse neurovirulence in EV-A71. Our data highlighted the key role of electrostatic interactions in shaping heparin-binding to confer virulence in mice. Among the weak heparin-binding variants used in this study (KEK, EEK and EEI), only EEI was associated with increased virulence and virus fitness *in vivo*. Strikingly, EQI, which should be a strong heparin binder, showed high virulence, and we showed that this was due to the mutation VP1-K244E, which conferred weak heparin-binding. This VP1-K244E mutation has been previously identified as a mouse virulence determinant [43, 44]. Increasing polymerase fidelity or impairing recombination of EQI restricts the emergence of mutations and abolishes *in vivo* neurovirulence, suggesting the importance of viral population diversity in EV-A71 pathogenesis, similar to reports on poliovirus [45, 46]. We showed that selection of individual adaptive mutations with roles in heparin-binding impacts viral pathogenicity.

Six EV-A71 variants were engineered with different amino acids at VP1-98, VP1-145 and VP2-149. VP1-98 and VP1-145 also have important roles in binding to PSGL-1 found in blood cells [8]. Both VP1-145 and VP1-149 have been implicated in mouse adaptation and virulence [22, 23, 25, 43, 47]. The VP1-98 and VP1-145 residues also act as modulators of heparin-binding in cell culture [36]. EV-A71 uses heparan sulfate as an attachment receptor, but it is not known how this drives pathogenesis. Heparin-binding may impact virulence outcome through neuroinvasion or neurovirulence [48]. Unlike SCARB2, heparin has no role in viral uncoating and internalization [7, 49–51], suggesting that heparin-dependent virulence is unlikely to be due to virus-functional receptor interaction. In mice, mSCARB2 and mPSGL-1 are known to poorly support EV-A71 infection [49, 52, 53]. Therefore, an unidentified mouse receptor could be utilized to achieve high viremia and dissemination. The cell binding properties of the engineered variants were as expected, with EEI, EEK and KEK showing lower binding due to VP1-145E and/or VP1-98E; and EQK, KQK and EQI showing higher binding due to VP1-145Q. VP1-149K and VP1-149I did not influence cell binding. This is probably related to mouse and human-specific tropism, as VP1-149I has only been reported in mouse studies [22, 43, 47].

Establishment of viremia is crucial for further dissemination to other target tissues such as skin and invasion into CNS [54]. During *in vivo* dissemination, the strong binding affinity of a heparin binder increases the likelihood of virus being sequestered by tissue GAG, resulting in rapid virus clearance from blood circulation [31, 33, 55]. This gives rise to a low viremia level with a substantial reduction of virulence in mice. Kobayashi and colleagues reported that EGK virus is less virulent compared to EEK due to the former harbouring the VP1-G145 residue, enabling the virus to adsorb more strongly to HS, resulting in attenuated virulence in SCARB2-expressing transgenic mice [56]. Our findings that a strong heparin-binding phenotype attenuates virulence is also observed in other viruses, including Sindbis virus [33], Venezuelan equine encephalitis virus [55], West Nile virus [30], yellow fever virus [31] and Japanese encephalitis virus [57]. Using a monkey model, Zhang *et al.* showed that the establishment of viremia was strongly correlated with EV-A71 neuroinvasion into CNS [16]. A clinical study correlating prolonged viremia in EV-A71 patients with severe CNS involvement further suggests the importance of viremia in determining severity outcome [58].

We performed intracerebral infection to bypass peripheral dissemination and neuroinvasion, therefore directly measuring the neurovirulence of each variant. Unlike the weak heparin binders EEI and EQI-K244E^+^, intracerebrally inoculated EQI failed to exhibit the same neurovirulent phenotype as it did following i.p. administration, for which we propose two possible explanations. First, virus replication in brain cells may be restricted. This is supported by histopathological studies showing very few neurons in brain are infected with EV-A71 [59, 60]. EQI viruses failed to acquire the VP1-K244E mutation, presumably due to lower infectivity of neurons and suboptimal replication following direct inoculation into brain, resulting in low virulence in mice. Secondly, viral multiplication in extraneural tissues such as hind limb skeletal muscle plays a key role in neuropathogenesis [61]. Mice intracerebrally inoculated with EEI showed higher viral load in hind limb muscles (S1 Fig), implying that the virus spread to peripheral tissues and underwent further extraneural replication (especially in the skeletal muscles) before re-entering the brain at high titers [16, 25, 61]. Retrograde axonal transport is the main transmission route for neuroinvasion [60, 62]. High replication will result in muscle damage that increases retrograde axonal transport and virus trafficking to the CNS [63, 64]. The weak heparin binders EEI and EQI-K244E^+^ remain lethal since they disseminate effectively and establish high viremia prior to neuroinvasion. Using an *in vitro* porcine blood brain barrier (BBB) model (S1 text), we found no correlation between heparin-binding and neuroinvasion across the BBB through tight junction leakages, as demonstrated by EEI and EQI which induce poorer permeability compared to wild type EQK (S2 Fig). This suggests that hematogenous spread is not the main route of spread to brain [62]; rather, establishment of high viremia appears crucial for virus dissemination to other target organs which support high levels of replication.

The mechanism of pathogenesis associated with heparin-binding is driven by electrostatic interactions at the five-fold axis of the virus surface. Alteration of the five-fold axis may affect capsid instability resulting in conformational changes which trigger genome uncoating, bypassing the need for receptor-virus binding. In poliovirus, mouse adaptation is controlled by a balance between capsid plasticity during uncoating and thermostability of the virion [65]. Similarly, electrostatic repulsion around the five-fold axis which results in capsid instability is observed in naturally thermo-labile foot-and-mouth-disease virus [66]. In EV-A71, the observed dynamic switching between weak and strong heparin-binding phenotypes highlights the importance for the virus to maintain an optimal electrostatic interaction for stable capsid conformation. Multiple VP1-244 variants (K244, T244 and E244) were generated with different effects on electrostatic interactions with heparin (K>T>E). As K to E represents a non-conservative substitution, the virus has evolved to transition through a non-charged residue. (T) which has lower fitness cost during viral dissemination [67, 68]. As GAGs are ubiquitous in tissues, a natural selective pressure thus exists to revert weak heparin-binding variants to heparin-binding variants which can then attach to and infect a range of host cells.

Three complementary mutations, VP1-L97R, VP1-N104S and VP1-E167G, were detected along with VP1-K244E from EQI-infected moribund mice. Given that these mutations are near the VP1-244 residue site and located at loops (S3 Fig A), they could be selectively utilized by the virus to stabilize the conformational structure of the VP1-K244E mutation at the five-fold axis. VP1-N104S was frequently associated with the VP1-244 variants. Root mean square fluctuation (RMSF) was measured to examine the dynamic movement of residues (S3 Fig B); the RMSF value of the BC loop (where VP1-N104S is located) was relatively higher in EQI-K244E compared to other virus variants. This indicates that residues within the VP1 BC loop are flexible and bound poorly to heparin. The emergence of the VP1-N104S mutation could contribute to the conformational stability of VP1-K244E, resulting in enhanced viral infectivity [69]. The VP1-L97R mutation, within the VP1 BC loop, has been reported to enhance heparin-binding [70]. This mutation was first detected from EV-A71 in the blood, CSF and stool from an immunocompromised patient [71]. However, we found this mutation in the skeletal muscle but not in the brain of a moribund mouse. Similarly, the presence of the VP1-L97R could stabilize the conformational structure of VP1-K244E. Interestingly, both VP1-L97R and N104S did not co-exist in the same sample. The VP1-D31G [72] and VP1-E167G [71] mutations previously implicated in neurotropism in humans were also observed in the brain samples but at very low frequencies. Dynamic switching between weak and strong heparin-binding phenotypes has been observed *in vitro* [36, 73]. Furthermore, restricting polymerase fidelity and impairing recombination rendered the virus avirulent, suggesting that restriction of population diversity alters fitness of the virus *in vivo*. In the present study, only VP1 was sequenced, and there may be other genomic changes within the virus genome.

We propose that our findings are not only limited to the murine model but may reflect neuropathogenesis in humans. Strong heparin-binding variants (EQK and EGK) have been more frequently detected from sequencing of virus cultures than from direct sequencing of clinical specimens, suggesting that heparin-binding phenotypes are a consequence of adaptation to tissue culture [18, 20, 23, 71, 74–101] (S1 Table). Detection of VP1-E145 in the sequences of a fatal encephalitis autopsy specimen [102] further reinforces our view that weak heparin-binding is associated with virulence in humans, as we have shown in mice in the present study. EV-A71 infection is usually mild and limited to HFMD, with neurological complications seen in 0.1-1.1% and deaths in 0.1% to 0.03% [103–107]. The frequent reversion between heparin-binding and weak heparin-binding variants fulfils the trade-off hypothesis in which the virus juggles between the cost and benefits of harming the host. Nevertheless in humans, selection of weak heparin-binding viruses to cause severe neurological disease appears rare.

We have developed a hypothetical EV-A71 pathogenesis model to show the importance of heparin-binding in human infection (Fig 9). Three determinant factors of EV-A71 virulence are virus entry, dissemination and neuroinvasion. Both strong and weak heparin binders infect humans at the same rate. Primary viremia is established upon virus entry in primary replication sites such as tonsils and oropharynx [15]. Strong heparin binders are readily removed from blood circulation due to their high affinity to heparin, thereby reducing the viremia level. Weak heparin binders are not adsorbed into tissues, but remain in the bloodstream, undergoing further extraneural replication in skeletal muscles, giving rise to high viremia. Upon overcoming the immune system, the high viremia results in better replication and dissemination to skeletal muscles, and peripheral motor nerves, through which the virus invades the CNS by retrograde axonal transport [15, 108]. High viremia alone does not result in direct CNS invasion as the virus cannot traverse the BBB. The dynamic reversion between heparin-binding and weak heparin-binding phenotypes in viruses occurs due to the differing needs of the virus to bind to the highly-abundant GAGs in tissues and to disseminate widely. Many research questions however remain. What additional host factors could alter the heparin-binding phenotypes? Could the host immune responses influence the fate of the viruses if activated early enough before the high viremia stage?

**Fig 9.**
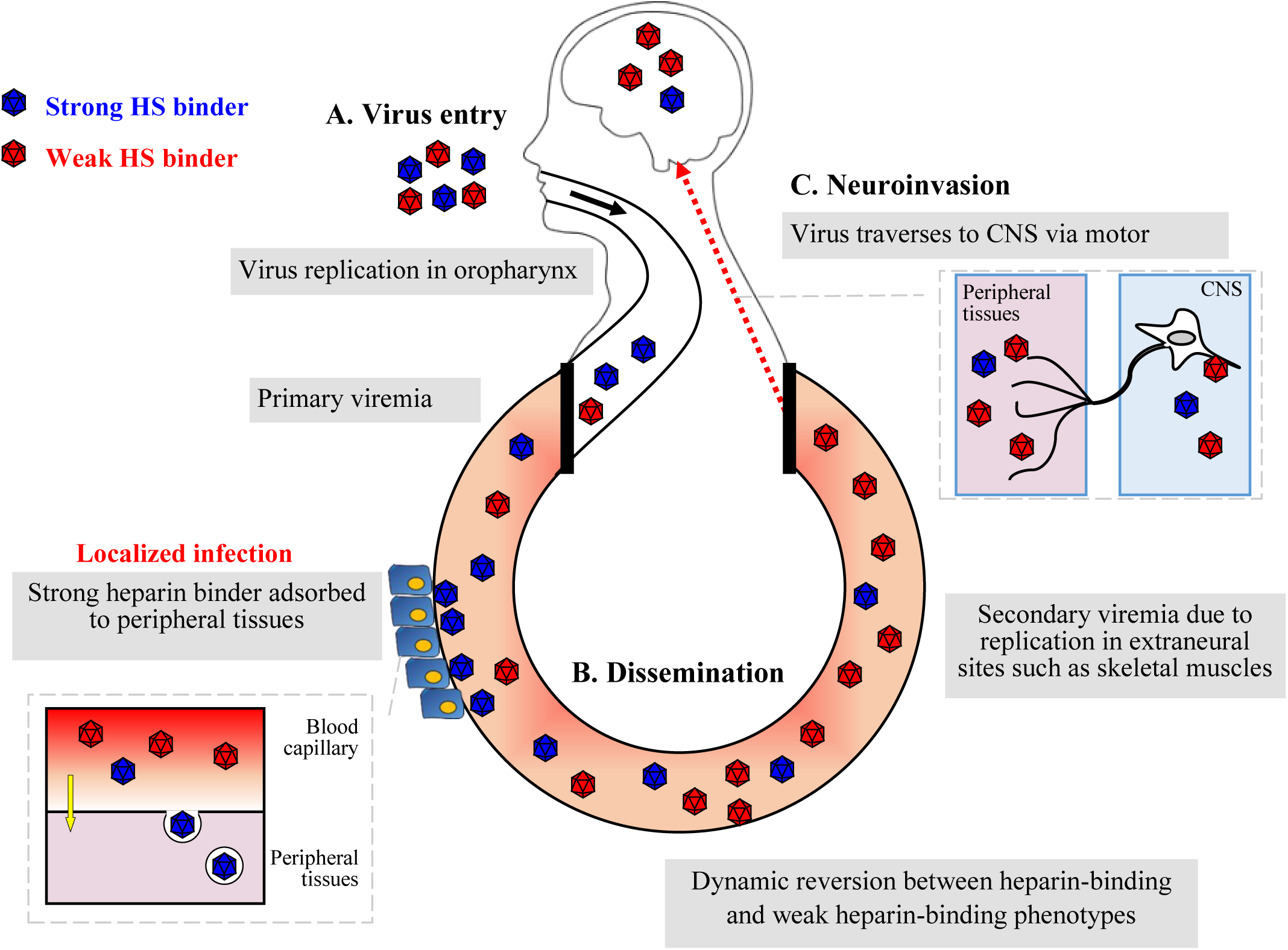
Hypothesized model of EV-A71 heparin-dependent pathogenesis in human. Three major factors are responsible for EV-A71 virulence determination, namely virus entry, peripheral dissemination and neuroinvasion. (A) Both strong and weak heparin binders infect humans at the same rate, using the same inoculation route and receptor. (B) Viremia is established upon virus entry. Strong heparin binders are more readily removed from the blood circulation by binding to peripheral tissues due to their high affinity to heparin. Meanwhile, weak heparin binders give rise to higher viremia with better dissemination to other organs. (C) Neuroinvasion occurs when virus travels from peripheral motor nerves to the CNS via retrograde axonal transport.

In summary, we showed that weak heparin-binding EV-A71 is highly virulent in mice, in contrast with strong heparin binders which show higher replication *in vitro* due to culture adaptation. This study shows that weak heparin-binding EV-A71 is preferentially selected to disseminate via the bloodstream; in contrast, strong heparin-binding EV-A71 is adsorbed to peripheral tissues and rapidly cleared. The electrostatic surface charges at the VP1 capsid shape heparin-binding and hence EV-A71 virulence. Our findings provide the mechanistic action of heparin-dependent virulence, and have potential therapeutic implications for viruses which utilize heparin as an attachment receptor and are dependent on high viremia levels to cause infection.

## Materials and Methods

### Ethics statement

The animal experiments were carried out in accordance with the rules and guidelines of the Animal Experimental Unit (AEU) in University of Malaya. The protocols were reviewed and approved by the Institutional Animal Care and Use Committee of the Faculty of Medicine, University of Malaya (reference number: 2016-190908/R/TCW).

### Cell lines and viruses

Human rhabdomyosarcoma (RD, ATCC no.: CCL-136) and human glioblastoma cells (U-87MG, ATCC no.: HTB-14) were propagated in Dulbecco’s Modified Eagle’s Medium (DMEM) (Life Technologies). Chinese hamster ovary cells (CHO-K1; ATCC no: CCL-61) were maintained in Kaighn’s modification of Ham’s F-12K nutrient mixture (Life Technologies). All cells were supplemented with 10% fetal bovine serum (FBS). Infected cells were maintained in media containing 2% FBS. All cells were maintained at 37°C in 5% CO_2_.

EV-A71 strain 41 (5865/SIN/000009, GenBank accession no. AF316321; subgenogroup B4) was used for construction of infectious clones using a DNA-launched strategy as reported previously [109]. Different mutations were incorporated into the EV-A71 infectious clone plasmid using Q5 high-fidelity DNA polymerase (NEB) PCR site-directed mutagenesis with primers listed in Table S2. The purified PCR products were treated with T4 polynucleotide kinase, T4 ligase and *Dpn*I (NEB) for 1 hour at room temperature. The ligation mixture was then transformed into *E. coli* XL-10 GOLD ultracompetent cells (Agilent Technologies). The plasmids were transfected into RD cells using Lipofectamine LTX (life Technologies) as reported previously [36].

### Evaluation of virus attachment in cell lines

The binding efficiency of EV-A71 variants to RD, U-87MG and CHO-K1 cells were evaluated using cell-based ELISA. Each virus variant was inoculated in triplicate wells in different cell lines at MOI of 20 (in 100 µl), and incubated for 1 hour at 4°C. Unbound viruses were washed away three times using cold PBS. The infected cells were then fixed with 4% paraformaldehyde for 20 minutes at room temperature. Antibody staining was performed using EV-A71 monoclonal antibody MAB979 (Merck, USA; 1:2500 dilution in 1 % BSA) for 1 hour at 37°C. After three washes with PBS-T (0.05% Tween 20 in PBS), HRP-conjugated anti-mouse IgG antibody (Gene Tex, USA; 1:2000 dilution in 1% BSA) was added to each well for 1 hour at 37°C. After a final wash, KPL TrueBlue peroxidase substrate (SeraCare, USA; 100 µl/well) was added. After 10 min, the reaction was immediately stopped by adding in stop solution and the absorbance reading was measured at 450 nm.

### Binding of EV-A71 particles to immobilized heparin sepharose beads

A binding assay was performed using columns with immobilized heparin sepharose beads as previously reported [36]. In brief, 200 µl of heparin sepharose (Abcam, UK) was aliquoted into a Pierce Spin Cup with cellulose acetate filter (Thermo Scientific, USA). The heparin sepharose beads were washed twice with binding buffer (0.02 M Tris-HCl, 0.14 M NaCl, pH 7.4), before addition of each virus variant (1 × 10^5^ PFU in 600 µl). The columns containing viruses were incubated for 30 minutes at 4°C, and this was followed by centrifugation and 5 washing steps. The heparin-bound viruses were collected after eluted with elution buffer (0.02 M Tris-HCl, 2M NaCl, pH 7.4). Both virus input and output fractions were quantitated using real-time PCR, and the virus binding efficiency was normalized by dividing the output viral RNA copies number over the input viral copies number.

### Evaluation of inhibitory effect of soluble heparin on EV-A71 variants

To determine the inhibitory effect of soluble heparin on EV-A71 variants, a virus inactivation assay was performed as previously described [9]. In brief, viruses were incubated with 2.5 mg/ml of soluble heparin (Sigma, USA) for an hour at 37°C. The treated viruses were inoculated onto pre-seeded RD cells and incubated at 37°C. Two days later, the cell viability of each infected virus variants was measured using CellTiter 96 Aqueous One solution Cell Proliferation Assay (Promega, USA). The relative cell viability was calculated by normalizing the absorbance value of treated virus samples against untreated virus samples, as compared to the mock-infected samples.

### Mice infection experiments

Groups of one-day old ICR suckling mice (n= 9 to 12) were obtained from AEU. Each group of suckling mice were either intraperitoneally or intracerebrally inoculated with 1 × 10^5^ PFU of each EV-A71 variant or PBS alone. All infected mice were monitored daily for weight change and health status up to 13 days post-infection. A clinical score was recorded using the following grades: 0, healthy; 1, weak or less active; 2, hunched posture and lethargy; 3, one-limb paralysis; 4, two-limb paralysis; 5, moribund or dead. Moribund mice were sacrificed and removed along with any mice found dead. Harvested mice organs were homogenized using hard tissue homogenizing mix (Omni International, USA). RNA was extracted from the homogenates with QIAamp viral RNA mini kit (Qiagen, Denmark). The viral loads in organs were determined using TaqMan fast virus 1-step master mix (ABI, USA). One step RT-PCR was also performed to amplify viral RNA from the organs using MyTaq One-Step RT-PCR kit (Bioline, UK) for Sanger sequencing or deep sequencing. Illumina Miseq (Illumina, USA) next-generation sequencing (NGS) was performed with 150 nucleotide paired end reads and average coverage of at least 20,000 reads. The NGS reads were analyzed using CLC Bio Genomic Workbench (Qiagen).

For the virus clearance assay, 3- to 4 week old ICR mice (weighing within 25-35g) were i.p. injected with ketamine/xylazine cocktail prior to infection. Anesthetized animals were then intravenously inoculated with 5 × 10^5^ PFU of each EV-A71 variant via the tail vein. At certain timepoints, blood was collected from anaesthetized mice through the retro-orbital plexus with the use of a sodium heparinized hematocrit capillary (Hirsschmann, Germany). The collected whole blood was then used for viral RNA quantitation.

### Immunohistochemistry

Immunohistochemistry (IHC) were performed by the standard ENVISION technique as described previously [110]. Briefly, deparaffinised and rehydrated tissue sections were blocked using standard immunoperoxidase procedure before antigen retrieval (30 minutes, 99°C, Tris EDTA buffer with 0.05% Tween-20). Tissues were then incubated with rabbit polyclonal EV-A71 VP1 (GeneTex, USA) at 4°C overnight. After washing, tissues were then incubated with goat-anti rabbit HRP-conjugate (Dako, Denmark) for 30 minutes at room temperature. Tissues were stained using DAB (Dako) and counterstained with hematoxylin (Dako). The tissues were mounted using DPX (Dako) prior to examination under a light microscope. The negative control tissues for IHC included mock-infected ICR mice brain and hind-limb muscle tissues. Isotype control antibodies or normal rabbit immunoglobulin fractions (Dako) were also used to exclude non-specific staining.

### Electrostatic surface charge analysis of EV-A71 structure

The EV-A71 structure was visualized using Chimera software (UCSF Chimera version 1.13.1, USA). Electrostatic surface potentials of virus capsid were analyzed using the ‘Coulombic surface coloring’ function in which the capsid residues were labelled with different colors based on their electrostatic charges. Positively-charged residues were colored blue while negatively-charged residues were colored red.

### Molecular docking simulation of EV-A71 VP1 and 12-mer heparin

Molecular docking simulation of EV-A71 crystal structure (PDB ID: 4AED) was performed using CDOCKER (CHARMm-based DOCKER) [111], as previously described [36].

### Statistics

All experiments were performed with at least two biological duplicates. Data are shown with error bars indicating standard deviations. Student’s *t*-test was performed for all *in vitro* experiments as well as viral load quantitation from mice organs. Survival of mice was evaluated using Kaplan-Meier analysis. GraphPad Prism version 5.03 (GraphPad Software, USA) was used for statistical analyses with a *P* value of < 0.05 indicating significance.

---

**S1 Table.** Comparison of EV-A71 isolate sequences of primary specimens and passaged isolates. Strong heparin binders (denoted with asterisks) were more frequently identified from sequencing of passaged EV-A71 than from direct sequencing of primary specimens, suggesting that the virus isolates have undergone heparin-binding adaptation in tissue culture (*P* < 0.00001, chi-square test).

| Sequence combination | Sequencing approach |  |  |  |
| --- | --- | --- | --- | --- |
|  | Direct sequencing |  | Sequencing from tissue culture propagation |  |
|  | Number of sequences | Percentage (%) | Number of sequences | Percentage (%) |
| EEK | 218 | 88.62 | 142 | 63.96 |
| EQK* | 4 | 1.63 | 13 | 5.86 |
| EGK* | 9 | 3.66 | 30 | 13.52 |
| KEK* | 13 | 5.28 | 31 | 13.96 |
| Others | 2 | 0.81 | 6 | 2.70 |
| Total | 246 | 100.00 | 222 | 100.00 |
| Chi-square test | $\chi^2 = 40.3558$. $P < 0.00001$ | | | |

**S2 Table.**
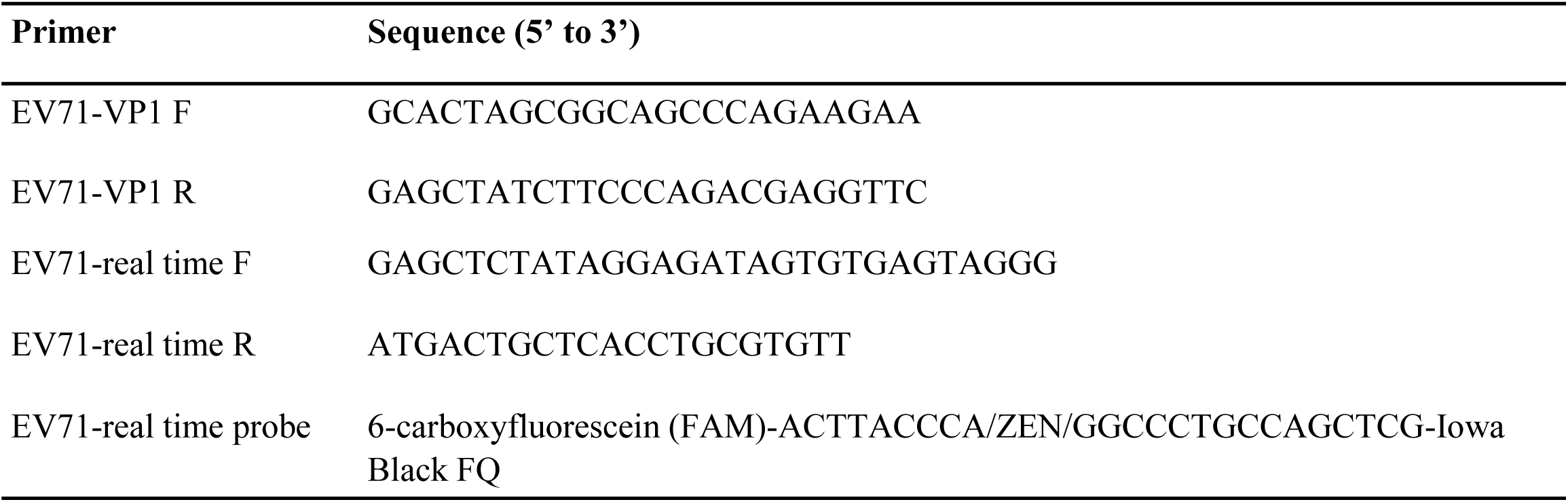
Primer sequences used for RT-PCR and qRT-PCR. Primer sets used for EV-A71 VP1 sequencing and qRT-PCR are shown.

**S1 Fig.**
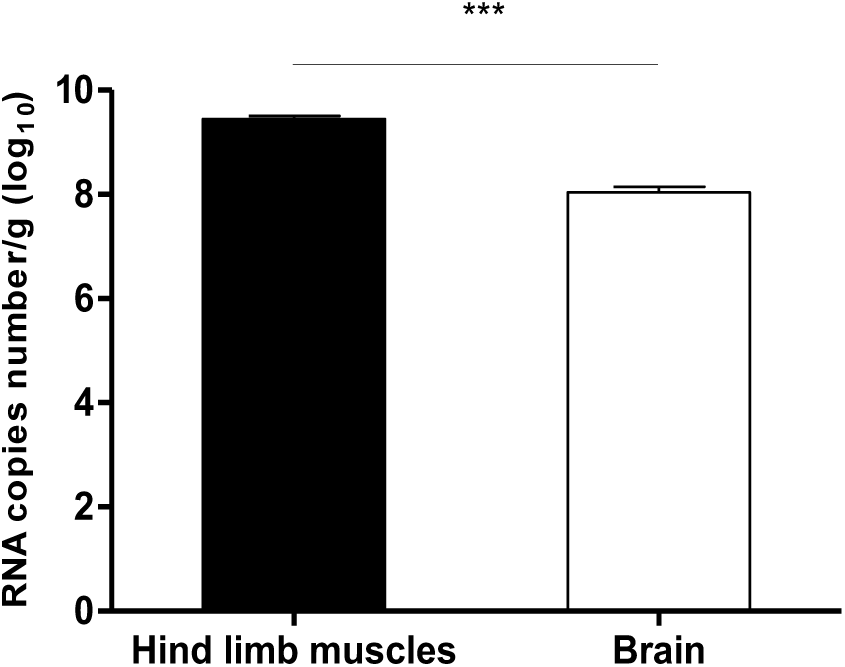
Comparison of viral loads from muscles and brain in mice infected through the i.c. route. One-day old suckling mice (n=3) were infected with EEI through i.c. route of administration. At day 4 post-infection, muscles and brains were harvested and viral loads were quantitated using qRT-PCR. Significant comparisons are labelled *** (*P* < 0.001).

**S2 Fig.**
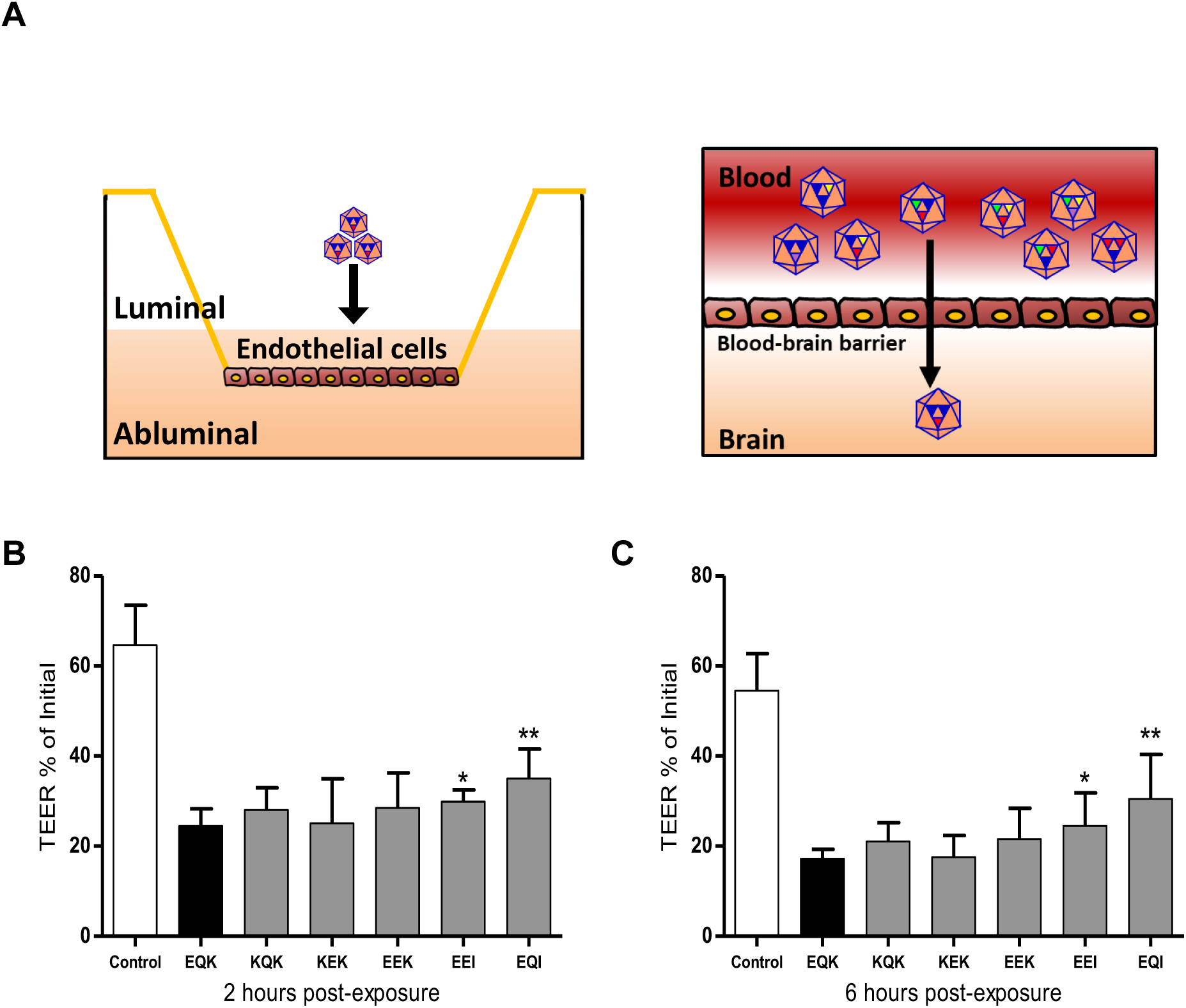
Effects of EV-A71 exposure on a porcine *in vitro* BBB model. Illustration of the porcine *in vitro* BBB model which simulates the movement of virus particles through an *in vivo* BBB, in which the luminal side represents the blood capillary while the abluminal compartment represents the brain (A). The *in vitro* model was exposed to different EV-A71 variants with titer of 1 × 10^5^ PFU. The BBB permeability induced by EV-A71 variants were assessed in terms of transendothelial electrical resistance (TEER), with a greater reduction of TEER indicating greater permeability of BBB through tight junction leakages. The TEER was recorded at 2 hours (B) and 6 hours post-exposure (C) along with non-infected cell controls (white bars) and normalized with TEER values measured before virus exposure. Results are presented as mean ± SD (n=6). Significant differences between viral variants and WT (black bars) are labelled as * (*P* < 0.05) and **(*P* < 0.01), using the Student’s *t* test.

**S3 Fig.**
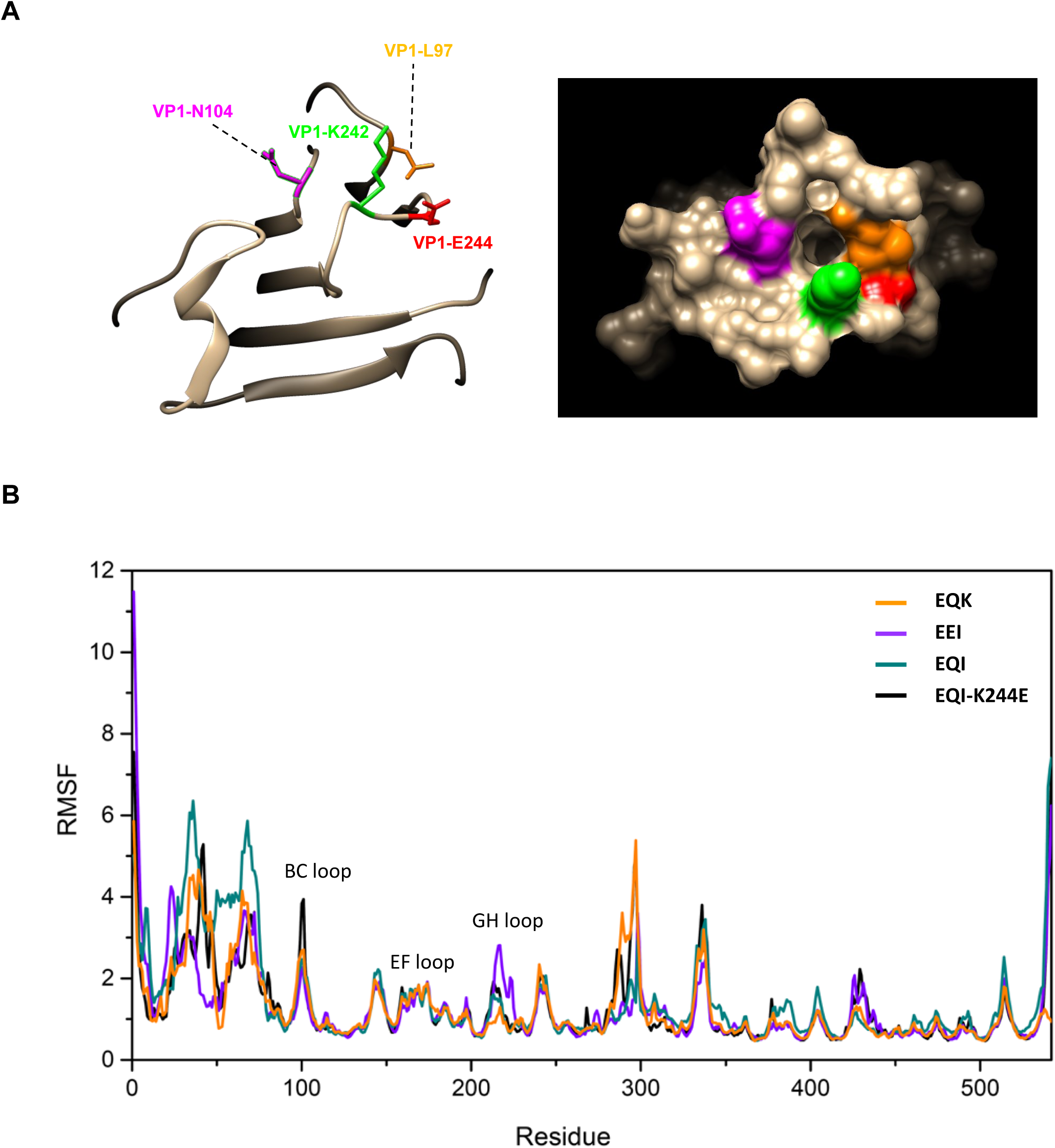
Structural modelling of EQI-K244E variant and root mean square fluctuation analysis of different EV-A71 variants. (A) Structural modelling of VP1 amino acid residues of EQI-K244E (left panel). Each important amino acid is labelled with different colors: VP1-E244 in red, VP1-K242 in green, VP1-L97 in orange and VP1-N104 in magenta. Note that VP1-E145 is not visible from this angle. The surface of EQI-K244E (right panel) is displayed corresponding to the structural model. (B) Root mean square fluctuation (RMSF) value of VP1 and VP2 amino acids are displayed for different variants. VP1 comprises residues 1-297 whereas VP2 consists of residues 298-542. BC, EF and GH loops of VP1 are labelled accordingly.

S1 Text. Establishment of *in vitro* blood-brain barrier model

Primary porcine brain endothelial cells (PBECs) were cultured as reported previously with slight modifications [112, 113]. To set up *in vitro* blood-brain barrier (BBB) model, the PBECs were seeded onto Corning Transwell inserts at a density of 1 × 10^5^ cells/cm^2^. Upon cell confluency, the culture medium was changed to a serum-free medium containing hydrocortisone (550 mM). The cells were treated with CPT-cAMP (250 μM) and RO-20-1724 (17.5 μM) for approximately 24 hours to induce BBB differentiation, followed by measurement of transendothelial electrical resistance (TEER).

TEER of the PBEC monolayer was measured using a STX-100C chopstick electrode pair connected to an EVOM meter (World Precision Instruments Inc., Sarasota) as an indicator for BBB tight junction function. The first measurement was conducted approximately 24 hours post CPT-cAMP and RO-20-1724 treatment, and prior to virus exposure. TEER of a blank filter insert without cells was subtracted from measured TEER of cell monolayers, and the values multiplied by surface area of the filter insert (1.12 cm^2^) to give the final unit of Ω.cm2. All monolayers used for this study showed TEER above 100 Ω.cm^2^.

EV-A71 variants were added to the luminal (‘blood-facing’) compartment i.e. the filter inserts, and the filter inserts was incubated at 37°C. After 1-hour incubation, medium containing virus samples was removed and replaced with fresh serum-free medium with added hydrocortisone (550 nM), and the inserts were returned to the incubator. Subsequently, TEER was measured at 2 and 6 hour post-exposure. At each time point, samples were aliquoted from the luminal and abluminal (‘brain-facing’) compartments for qRT-PCR quantitation. Mock infected controls were included for comparison.

